# Intermolecular proteolytic processing of SPRING and Site-1-protease regulate SREBP signaling

**DOI:** 10.1101/2023.03.28.534447

**Authors:** Sebastian Hendrix, Josephine M.E. Tan, Klevis Ndoj, Masoud Valiloo, Lobke F. Zijlstra, Roelof Ottenhoff, Nabil G. Seidah, Anke Loregger, Noam Zelcer

## Abstract

The SREBP transcription factors are central regulators of fatty acid and cholesterol metabolism. Produced as membrane-resident precursor proteins in the ER, their transcriptional activation requires the cholesterol-dependent translocation to the Golgi, and subsequent proteolytic cleavage by S1P, a type-I transmembrane protein. S1P is produced as a proprotein convertase that needs to undergo autocatalytic cleavage to attain its mature form in the Golgi, in a process that is not fully elucidated. We have recently identified SPRING (C12ORF49) as a novel regulator of the SREBP pathway and reported that S1P activity and retrograde recycling of the SREBP chaperone SCAP are affected. Here, we demonstrate that SPRING and S1P interact and that in co-transfection experiments in mammalian cells this facilitates the autocatalytic activation of S1P_A→C_ form. Accordingly, S1P_A→C_ processing of stably overexpressed S1P in SPRING^KO^ cells is attenuated, but not abolished, and does not rescue SREBP signaling. Reciprocally, we identified a conserved S1P cleavage site in SPRING, and demonstrate that cleavage of SPRING results in secretion of the SPRING ectodomain. SPRING cleavage is S1P-specific and can be pharmacologically inhibited by S1P inhibitors or by mutating the S1P cleavage site. Functional analysis revealed that the SPRING ectodomain was sufficient to support S1P_A→C_ processing and SREBP signaling, but that SPRING cleavage is not a prerequisite for this. In conclusion, our study reveals a complex interplay between the proteolytic activation of S1P and SPRING yet suggests that this is not the primary mechanism underlying the role of SPRING in SREBP signaling.

## Introduction

Cells must tightly control their sterol and fatty acid homeostasis to ensure that their metabolic and growth demands are met. Accordingly, disturbed lipid metabolism is associated with many human diseases, including cancer, neurodegeneration, and coronary artery disease. The Sterol-regulatory element binding proteins (SREBPs) are transcription factors that govern all facets of fatty acid and cholesterol metabolism by regulating the expression of sterol-responsive genes (1,2). The three SREBP isoforms, SREBP1a, SREBP1c, and SREBP2, share structural similarity, but have a distinct tissue distribution (3-5). Also, the different SREBP isoforms control a distinct set of sterol-responsive genes, with SREBP1c primarily regulating genes implicated in fatty acid synthesis such as fatty acid synthase (*FASN*) and acetyl-CoA carboxylase (*ACC*) (4), and SREBP2 regulating genes linked to cholesterol synthesis and uptake, including those encoding for the rate-limiting enzymes in cholesterol biosynthesis, 3- hydroxy-3-methylglutaryl-coenzyme A reductase (*HMGCR*) and squalene epoxidase (*SQLE*), and the low-density lipoprotein receptor (*LDLR*) (5). The third isoform, SREBP1a, regulates both sterol- and fatty acid-associated genes.

SREBPs are produced in their precursor form as membrane-associated endoplasmic reticulum (ER)-resident proteins that contain an N-terminal basic helix-loop-helix leucine zipper domain and a regulatory carboxyterminal region (1). Their cholesterol-dependent activation has been extensively reviewed and involves their translocation from the ER to the Golgi where they are subject to sequential proteolytic cleavage by the proteases Site-1 protease (S1P, also known as SKI-1) and Site-2 protease (S2P; encoded by *MBTPS1* and *MBTPS2*, respectively)(6-8). The released N-terminal SREBP domain then translocates to the nucleus to induce expression of sterol-responsive target genes.

S1P is the 8^th^ member of the proprotein convertase family (9,10). This family is comprised of 9 serine proteases that show similarities to bacterial subtilisin and yeast kexin (11). S1P is synthesized as a 1052 amino-acid inactive proenzyme that undergoes post-translational proteolytic processing to attain its active form (9,10,12). In a first cleavage step during translation, the signal peptide is removed by a signal peptidase at site A. This is followed by two sequential autocatalytic cleavage events at site B’/B (**<u>R</u>**K**<u>V</u>**F^133^↓ **<u>R</u>**S**<u>L</u>**K^137^↓) followed by site C’/C (**<u>R</u>**R**<u>A</u>**S↓LSL^169^ … **<u>R</u>**R**<u>L</u>**L^186^ ↓). Cleavage at the B’/B site is performed in the ER in *cis*. Following trafficking of the partially cleaved intermediate to the Golgi cleavage of the C’/C site occurs, resulting in mature S1P (10,13). In its active form, S1P cleaves several proproteins at the C-terminus of the consensus motif RX(L,V,I)Z↓, whereby X can be any amino acid, excluding Pro or Cys and Z is any residue (preferably Leu) except Val, Pro, Cys, or Glu (11,14). Accordingly, S1P is implicated in the cleavage-dependent activation of, amongst others, several transcription factors (11,15). Other than SREBP1 and 2 (1,10), those include ER stress response related transcription factors like activating transcription factor 6 (ATF6) (16,17), cyclic AMP-responsive element binding proteins CREB3 (18), CREB3L1 (19), CREB4 (20), and CREB3L3 (21). Other recently identified targets are α/β-GlcNAc-1-phospho-transferase (GNTAB) (22), the kinase FAM20C (23), the (pro)renin receptor (24), and various arenavirus surface glycoproteins (25).

Using a set of haploid genetic screens performed in mammalian cells we have recently identified SPRING as a previously unrecognized determinant of SREBP activity in mammalian cells and mouse liver (26), a finding later confirmed by others (27,28). We reported that SPRING is required for adequate proteolytic processing of SREBPs and SREBP-dependent signaling. We have shown that absence of SPRING leads to reduced SCAP abundance and of

SCAP retrograde trafficking from the Golgi to the ER. In aggregate, this leads to functional SCAP deficiency and consequentially impaired SREBP signaling (26). The observed SCAP deficiency is reminiscent to that observed when S1P activity is pharmacologically or genetically inhibited (29). In line with this notion, we have reported that absence of SPRING moderately attenuated the S1P-dependent ATF6-mediated ER stress-response in tunicamycin-treated cells (26), an observation consistent with others reporting also an interaction between S1P and SPRING (27,30). Collectively, these results are consistent with a model in which SPRING regulates S1P activity upstream of proteolytic SREBP activation. In this study we mechanistically address this hypothesis in mammalian cells.

### Experimental procedures

#### Chemicals, cell culture and transient transfection

The reagents and chemicals used in this study are listed in Supplementary Table 1. The human cell lines HEK293T, HeLa, and HepG2 cells were obtained from ATCC and grown at 37°C and 5% CO2 in Dubecco’s Modified Eagle Medium (DMEM) (Gibco) supplemented with 10% Fetal Bovine Serum (FBS), penicillin (100 units/mL) and streptomycin (100 µg/mL). Hap1 control and Hap1-SPRINGKO cells were previously described (26). Wildtype and Spring(-/-) primary fibroblasts were isolated from Spring(fl/fl) mice (26). Briefly, an ear biopsy was obtained, digested with collagenase (1.3 mg/ml, Sigma-Adrich) followed by trypsinization (0.25% Trypsin-EDTA, Sigma-Aldrich) and passed through a 70 µm cell strainer to release the cells. Subsequently, cells were plated and grown in Roswell Park Memorial Institute 1640 (RPMI) (Gibco) supplemented with 10% Fetal Bovine Serum (FBS), 50 µM β-mercaptoethanol (Sigma-Aldrich), 100µM L-asparagine (Sigma-Aldrich), penicillin (100 units/mL) and streptomycin (100 µg/mL). The cells were immortalized by infection with a retrovirus encoding the Large-T antigen, and selected in medium containing 3 µg/mL puromycin. Following immortalization, L-asparagine and β-mercaptoethanol were removed from the culture medium. Immortalized cell populations were transfected with constructs encoding GFP or Cre-GFP (Addgene #49055 and #49056, respectively), and FACS sorted to yield single cell clones of Spring(-/-) and Spring(fl/fl) fibroblasts. Multiple independent clones were isolated and Spring expression and function was evaluated. Where indicated, to deplete cellular sterols cells were washed with PBS and culture media was replaced with RPMI containing 10% lipoprotein deficient serum, supplemented with 100 µM mevalonate (Sigma) and 5 µM simvastatin (Calbiochem), as previously reported (31). To block S1P activity the inhibitor PF429242 (Bio-Techne), dissolved in DMSO, was added to the culture media at a concentration of 10 µM unless stated otherwise. As control, in these experiments an equal volume of vehicle was added to the culture media. HeLa, HepG2 and mouse fibroblast cells were transiently transfected with the indicated expression constructs using JetPrime (Polyplus-transfection), according to manufacturer’s instructions. For transient transfection of HEK293T cells polycation polyethylenimine (PEI, Polysciences) was used. Equal transfection efficiency was monitored by co-transfection with an expression plasmid for GFP or mCherry, and was consistently comparable. To monitor secretion of SPRING, cells were transiently transfected with the indicated SPRING and/or S1P constructs. Where indicated, PF429242 was added to cells to inhibit S1P. Transfected cells were cultured for 48 h, cell lysates and culture medium were collected and immunoblotted for SPRING and S1P. All mouse experiments were approved by the Committee for Animal Welfare (University of Amsterdam).

#### Molecular cloning and generation of lentiviral particles for transduction of cell lines

The cloning of the human SPRING open reading frame and generation of corresponding expression constructs has been previously reported (26). Site directed mutagenesis was used to introduce the R45E mutations into pDONR221-SPRING-Myc. The resulting pDONR221-SPRING_R45E_-Myc plasmid was used to generate corresponding expression constructs following LR gateway recombination with pLentiCMV-Puro-DEST(670-1) (Addgene #17293). A similar strategy was followed to generate SPRING_1-44_ (SPRING stump), SPRING_45-205_ (secreted SPRING; sSPRING; preceded by the signal peptide sequence of PCSK9) and SPRING_R43E,R45E,L47E_ (uncleavable SPRING; unclSPRING). S1P-V5 was amplified from pIRES S1P-V5 EGFP, cloned into pDONR221 to generate pDONR221-S1P-V5, and LR recombined into pLentiCMV-Puro-DEST(670-1) as above (Addgene # 17293). To generate S1P-mCherry a 2-step assembly PCR was performed to combine the S1P CDS derived from pLentiCMV-Puro-DEST(670-1)-S1P-V5 and the mCherry CDS from pcDNA6.2-C-mCherry-DEST flanked by gateway cloning sites (A kind gift from Dr. Paul Beare, NIH). The resulting amplicon was introduced into pDONR221 yielding pDONR221-S1P-mCherry and from there LR recombined into pLenti6.3-DEST-V5 as above. pIRES-S1P-BTMD-EGFP and p3xFLAG CMV7.1-ATF6ɑ were previously reported (32). Correctness of all constructs used in this study was confirmed by digestion analysis and sequencing (Supplementary Table 2). To stably express *SPRING* and *S1P* constructs we transduced Hap1 and mouse fibroblasts with lentiviral particles encoding the corresponding constructs. Briefly, lentiviral particles were generated by transfecting HEK293T cells with the lentiviral expression construct together with 3^rd^ generation packaging plasmids, as previously reported (33). Cells were transduced for 16 hours using the supernatant containing the lentiviral particles at a 4:1 ratio with complete DMEM or RPMI medium, supplemented with 12 µg/mL polybrene (Santa Cruz). Cells were subsequently selected with puromycin at a concentration of 1.5 µg/mL and single clones isolated by limited dilution for subsequent studies. Cell viability was determined by using the MTT assay (Invitrogen) was used according to manufacturer’s instructions.

#### Immunoblot analysis and co-immunoprecipitation

Total cell lysates for immunoblotting were prepared in radio-immunoprecipitation assay (RIPA) buffer (Boston Biochem), which was supplemented with 1 mM phenylmethylsulfonyl fluoride (PMSF) (Sigma) and a protease inhibitors cocktail (Roche). Subsequently, lysates were cleared by centrifugation at 4°C at 12000 x g for 10 minutes. The protein concentration of the cleared lysates was determined using a BCA assay (ThermoFisher) following the manufacturer’s protocol, and an equal amount was loaded for analysis. Samples were separated on NuPAGE Novex 4-12% Bis-Tris gels (ThermoFisher) and transferred to nitrocellulose membranes, blocked in 5% milk (Elk) in PBS supplemented with 0.05 % Tween and subsequently probed with primary antibodies. For co-Immunoprecipitation (Co-IP) experiments, total cell lysates were prepared in NP-40 buffer (150 mM NaCl, 5 mM EDTA, 1% NP-40, 50 mM Tris-HCl, pH 7.4) supplemented with protease inhibitors (Roche) and phenylmethylsulfonyl fluoride (Sigma) and incubated overnight at 4 °C with the indicated antibodies. The protein-antibody complex was then captured with Protein G Dynabeads (Thermo Fisher) at room temperature for 1.5 h. Subsequently, beads were collected using a DynaMag magnet (Thermo Fisher) and washed three times with RIPA buffer. Bound proteins were eluted in NuPAGE sample buffer and analyzed by immunoblotting as indicated. The primary antibodies used in this study are listed in Supplementary Table 3. Secondary HRP-conjugated antibodies (A28177 & A27036, Invitrogen) were used and visualized with chemiluminescence on an IQ800 (GE Healthcare), and quantified using the ImageJ2 (Version 2.9.0/1.53t) gel analyzer plugin. Unless indicated, immunoblots shown are representative of at least 3 independent experiments with similar results.

#### Quantitative PCR

RNA was isolated from cells using the Direct-zol RNA miniprep kit (Zymo Research) following the manufacturer’s protocol. cDNA was generated using the iScript reverse transcription reagent (BioRad). SensiFAST SYBRgreen (Bioline) was used for real-time quantitative PCR (RT-qPCR) on a LightCycler 480 II system (Roche), and gene expression was normalized to the expression level of 36B4 and GAPDH. Data are presented as fold change calculated using the ΔΔCT method. Primer sequences are shown in Supplementary Table 4.

#### Immunofluorescence

Immortalized mouse Fibroblasts stably expressing S1P-mCherry were seeded on fibronectin (0.001% in PBS; Sigma Aldrich) coated cover slips. After overnight attachment, cells were washed two times with warm PBS and fixed using 4% paraformaldehyde (Sigma Aldrich F8775). Cells were permeabilized in 0.5% Triton X-100 (Sigma), aldehyde residues were quenched with 20mM Glycine and blocked with 2% BSA/PBS (Sigma). Next, cells were stained with a primary antibody against Golgin97 (1:200, Cell Signaling) for 1 h at room temperature. Followed by an 1 h incubation with a secondary chicken anti-rabbit antibody conjugated to Alexa-Fluor-488 (1:250 in 2% BSA/PBS; Thermo Fisher). Nuclei were stained for 20 min using 250 ng/mL DAPI in 2 % BSA/PBS (Thermo Fisher). Cells were washed 3 × 3 minutes in 0.5 % BSA followed by a final wash with H2O for 3 minutes and mounted on standard glass slides using Mowiol (Sigma). Imaging was performed on a Leica TCS SP8 X confocal microscope mounted on a Leica DMI6000 inverted microscope using LAS-X software. ImageJ2 software (Version 2.9.0/1.53t) was used to calculate the ratio of Golgi resident S1P-mCherry vs. total S1P-mCherry per cell. S1P-mCherry localization to Golgi was scored based on co-localization with the Golgin97 signal.

#### Statistics

Statistical significance was tested using ANOVA with Holm-Šidak post hoc analysis, multiple unpaired t-tests with Holm-Šidak post hoc analysis or t-test with Welch correction. When the assumption of normality was violated, Kruskal-Wallis with Dunn’s multiple comparison test was employed as an alternative to the ANOVA and Mann Whitney with Holm-Šidak correction for multiple comparisons. Outliers were identified using the ROUT analysis. Graphpad Prism version 9.0 was used for statistical analyses with a significance threshold of *p*<0.05. SEM or SD is indicated by error bars and *p* values are indicated by asterisks: \**p* < 0.05, \*\**p* < 0.01 \*\*\**p* < 0.001 and **** p < 0.0001

## Results

We recently found that absence of SPRING attenuates SREBP proteolytic activation in cells, resulting in reduced SREBP signaling (26). As also previously reported, this occurred in the absence of changes in S1P expression (Supplementary Figure 1). This is consistent with abrogated S1P activity. Hence, we reasoned that decreased S1P activity in the absence of SPRING will render cells more sensitive to S1P inhibition. Confirming this notion, we found that Hap1 cells or immortalized mouse fibroblasts lacking SPRING are indeed hypersensitive to the S1P inhibitor PF429242 (Figure 1A,B). Further supporting this notion, tunicamycin-induced cleavage of ATF6, another established S1P target (16), was reduced in *Spring*-devoid cells (Supplementary Figure 2) (26). To explore the underlying SPRING-dependent mechanism for reduced S1P activity we first tested whether these two proteins interact by co-IP. In these experiments we could demonstrate their bi-directional interaction (Figure 1C,D). These data show that both proS1P and its mature C-form lacking the pro-domain can interact with SPRING, possibly implicating the catalytic subunit of S1P in the binding to SPRING. We then aimed to fine map the domains mediating this interaction. We used the luminal domains of both SPRING (SPRING_49-205_, sSPRING), the region containing the cysteine-rich region, and that of S1P (S1P before transmembrane domain, S1P-BTMD). Both proteins are retrieved in the culture media, as anticipated, and interact as assessed by co-immunoprecipitation (Figure 1E,F).

**Figure 1.**
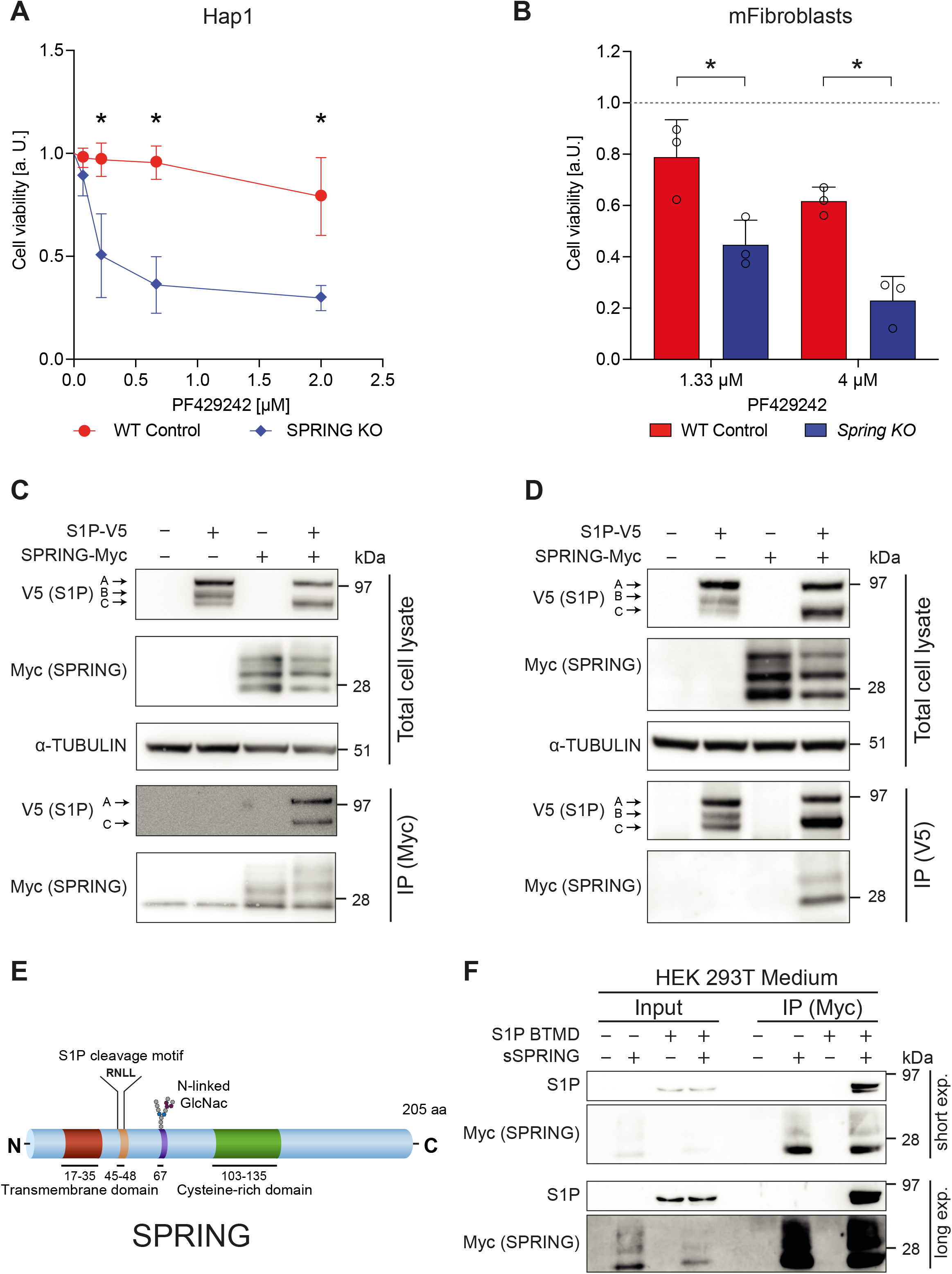
SPRING and S1P interact. Control or SPRING^KO^ (***A***) Hap1 cells or (***B***) immortalized mouse fibroblasts were cultured in the presence of the indicated concentration of the S1P inhibitor PF429242. Cell viability was evaluated with the MTT assay after 3 days (N=3). Dashed line indicates control cells grown in the presence of vehicle. (***C***,***D***) HEK293T cells were transiently transfected with SPRING and S1P expression constructs as shown. Total cell lysates were immunoblotted or used for co-immunoprecipitation as indicated (N=3). *Note:* A,B,C indicates the different S1P proteolytic intermediates. (***E***) Schematic representation of the domain structure of SPRING, with the predicted S1P cleavage site highlighted. (***F***) HEK293T cells were transiently transfected with the indicated expression constructs for secreted S1P (S1P before transmembrane domain; S1P-BTMD) and secreted SPRING (SPRING_49-205 ;_ sSPRING). Culture medium was used for co-immunoprecipitation and immunoblotted as indicated (N=3). Long and short exposures are shown. All immunoblots are representative of at least 3 independent experiments and bars and errors represent mean ± SD; * *p* < 0.05.

For activation S1P must undergo a complex set of auto-proteolytic events, involving the cleavage of the B’\B and C’\C sites and subsequent release of the pro-domain (Figure 2A) (9,10). In our initial experiments we observed a marked change in the pattern of the identified S1P bands when S1P was co-expressed with SPRING (Figure 1C,D). Co-expression of SPRING prompted a substantial decrease in the S1P_B_/S1P_C_ ratio and hence enhanced S1P activation to its C-form (Figure 2B). We also attempted to compare processing of endogenous S1P in wildtype and SPRING^KO^ cells, but lack of reliable commercial anti-S1P antibodies precluded this. Remarkably, while SPRING enhanced S1P processing into its mature C-form, S1P in turn prompted a reduction in abundance of full-length SPRING (Figure 2C,D). This was associated with detection of a higher mobility SPRING fragment in the culture medium (Figure 2D). The same was also observed when the experiments were repeated in HeLa and HepG2 cells (Supplementary Figure 3 and not shown). This finding points towards SPRING itself being subject to S1P-mediated cleavage, in line with the presence of a predicted RXLZ↓ cleavage-motif common to other S1P substrates (Figure 1E). Indeed, pharmacological inhibition of S1P with PF429242 or introducing the motif-disturbing mutations in the predicted cleavage site (SPRING_R45E_ or SPRING_(R43E,R45E,L47E)_ ; uncl. SPRING) abolished cleavage of SPRING in the presence of S1P (Figure 2D,E and Supplementary Figure 4). No high-resolution structure of SPRING has been reported to date. However, analysis of the Alphafold predicted structure positions this S1P cleavage site at the end of a flexible α-helical strand, preceding the luminal cysteine-rich domain, a position common to other S1P substrates (Figure 2F) (34). Collectively, this set of experiments reveals a complex interaction between SPRING and S1P that results in their reciprocal proteolytic processing and enhanced activation of S1P via the more efficient generation of its C-form that is devoid of the inhibitory pro-domain.

To test the functional significance of this proteolytic tango we first evaluated whether SPRING cleavage is a prerequisite for SREBP signaling. We stably reconstituted wildtype, and cleavage-mutant SPRING_R45E_ in Hap1- and mouse fibroblast SPRING^KO^ cells and tested their transcriptional response to sterol depletion. As previously reported (26), absence of SPRING results in a severely impaired SREBP-dependent transcriptional response to sterol depletion (Figure 3A,B). In this setting, the cleavage-mutant SPRING_R45E_ restored the sterol-dependent transcriptional response. Intriguingly, reintroducing SPRING_49-205_ that corresponds to the S1P-generated cleaved secreted SPRING fragment (sSPRING) was also able to restore SREBP-dependent signaling to levels comparable to those in control cells. In contrast, introducing the N-terminal SPRING stump (SPRING_1-44_) that is generated by S1P cleavage is unable to do so (Supplementary Figure 5). Rescue of the transcriptional response was also mirrored by a corresponding increase in the levels of the encoded proteins; Introducing sSPRING (SPRING_49-205_) restored SREBP signaling in Hap1 SPRING^KO^ cells (Figure 3C). Rescue of SREBP signaling was also seen when we introduced the cleavage mutants, SPRING_R45E_ and SPRING_(R43E,R45E,L47E)_ in Hap1 and mouse fibroblast SPRING^KO^ cells, respectively (Figure 3D,E). We also tested whether conditioned culture media from cells overexpressing SPRING_49-205_ can induce SREBP-dependent signaling in HepG2 cells, but in initial experiments were unable to see an effect of this treatment.

**Figure 2.**
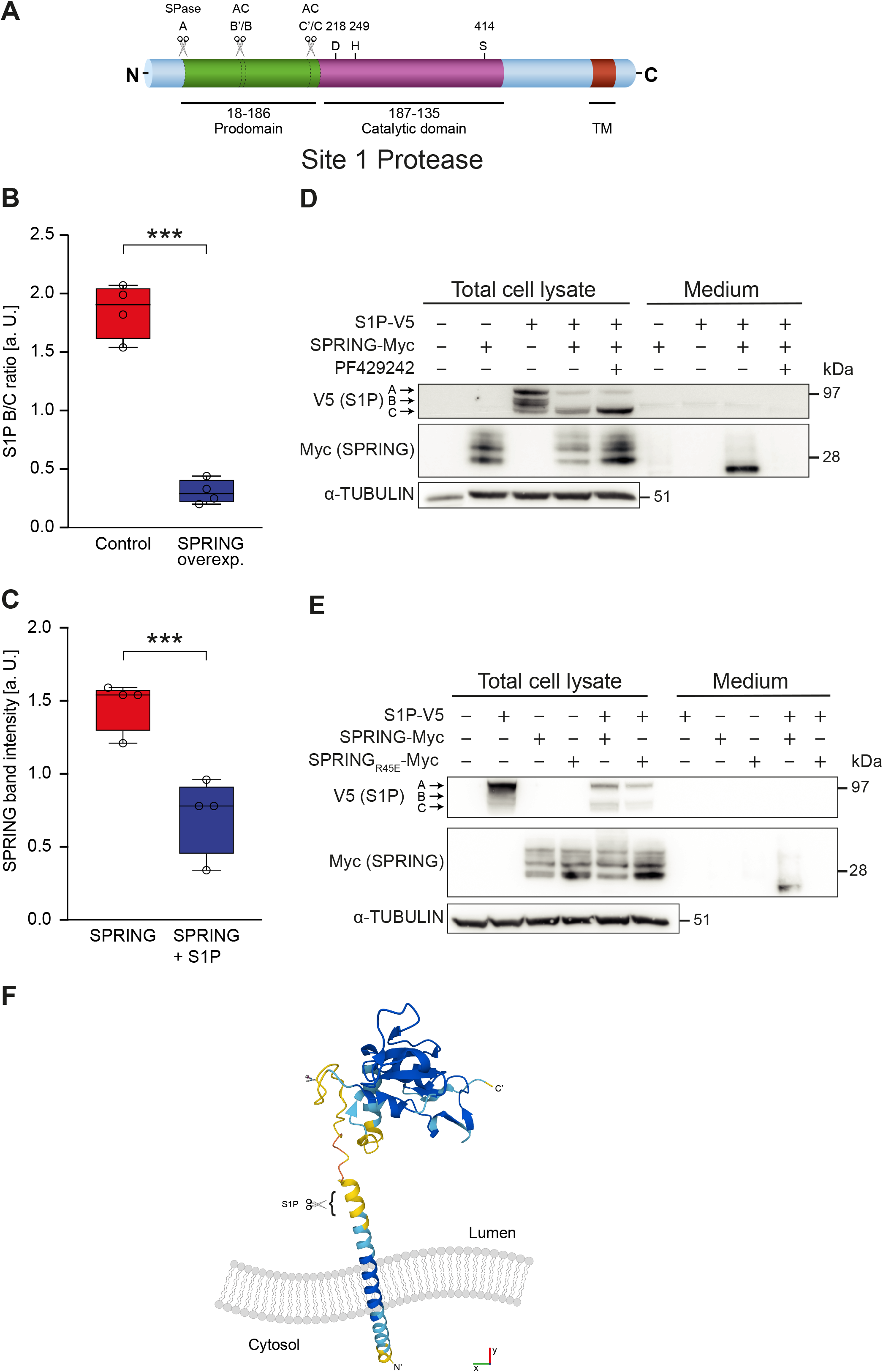
Bi-directional intermolecular proteolytic processing of S1P and SPRING. (***A***) Schematic representation of S1P domain structure. The A,B, and C proteolytic sites in S1P are highlighted. (***B***) Intensity of the S1P_B_ and S1P_C_ bands was quantified in cells transiently transfected with S1P and SPRING expression constructs and the S1P_B_/S1P_C_ ratio is shown (N=4). (***C***) Intensity of all SPRING bands was quantified in cells transiently transfected with S1P and SPRING expression constructs and the relative band intensities are shown (N=4). (***D***,***E***) HEK293T cells were transfected with the indicated S1P and SPRING expression constructs. Where indicated, cells were treated with 10µM of the S1P inhibitor PF429242 for 16 hrs. Total cell lysates and culture medium were immunoblotted as indicated. (***F***) The structure of SPRING as predicted by Alphafold. The predicted S1P cleavage site is highlighted. All immunoblots are representative of at least 3 independent experiments and bars and errors represent mean ± SD; *** *p* < 0.001.

**Figure 3.**
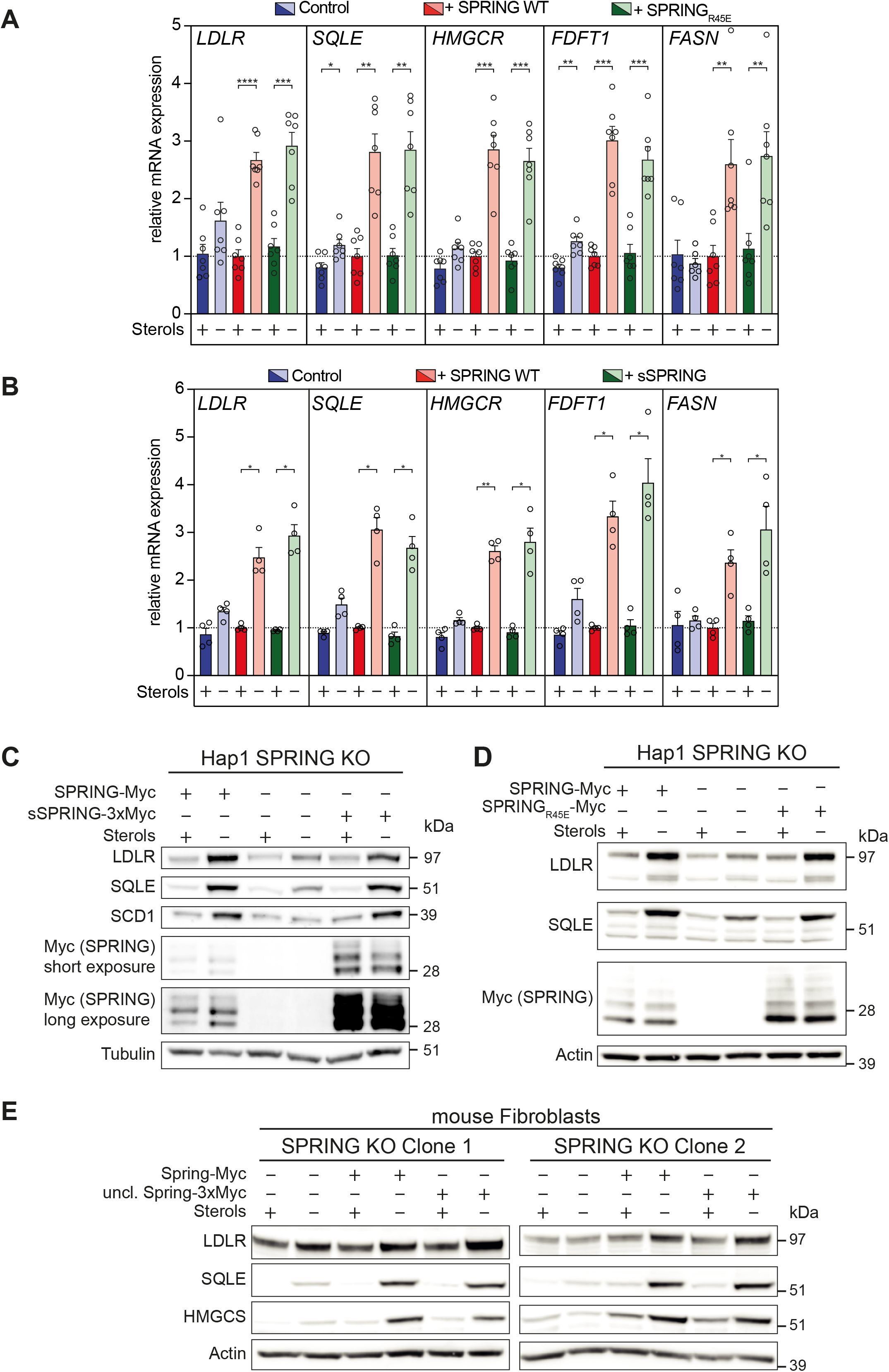
Cleavage-resistant and secreted SPRING support SREBP signaling in Hap1 cells and immortalized mouse fibroblasts lacking *SPRING*. (***A***,***B***) Hap1-SPRING^KO^ cells were transduced with the indicated SPRING constructs. Stable cells expressing wildtype wildtype SPRING, SPRING_R45E_, sSPRING (SPRING_49-205_), or control were grown in complete (+ sterols) or in sterol-depletion (-Sterols) culture media for 16 hrs. Expression of the indicated genes was determined by qPCR (N=4-7). (***C***,***D***) Cells were grown as in (***A***,***B***) and total cell lysates were immunoblotted as indicated. (***E***) Immortalized mouse fibroblasts lacking *Spring* expressing wildtype SPRING, SPRING_(R43E,R45,L47)_ (uncl. SPRING) constructs were grown in complete (+ sterols) or in sterol-depletion (-sterols) culture media for 16 hrs. Total cell lysates from two independent clones were immunoblotted as indicated. All immunoblots are representative of at least 3 independent experiments and bars and errors represent mean ± SEM; * *p* < 0.05, *** *p* < 0.001.

Having ruled out S1P-mediated proteolytic cleavage of SPRING as a prerequisite for maximal SREBP-dependent signaling in response to sterol depletion, we moved to consider impaired S1P processing as the primary lesion in cells lacking SPRING. We generated cells that stably overproduce S1P-V5 or S1P-mCherry and evaluated SREBP signaling in response to sterol depletion, as above. Consistent with the initial co-transfection experiments (Figure 1), absence of SPRING mildly effected processing of stably expressed S1P, as reflected by a significant increase in the S1P_B’/B_/S1P_C’/C_ ratio (Figure 4A,B). No signal was detected in control cells (*i*.*e*. non S1P-mCherry expressing) (Supplementary Figure 6). In these experiments the S1P_C’_ isoform was readily detected in both wildtype and *Spring*^*KO*^ fibroblasts, indicating that maturation of S1P is not dependent on SPRING and can take place in its absence. The same was the case when introducing S1P-V5 into Hap1-SPRING^KO^ cells (Figure 4C). Yet remarkably, despite intact processing of S1P, the SREBP-dependent transcriptional response remained refractory to sterol depletion in both fibroblasts- and Hap1*-*Spring^KO^ cells (Figure 4A,C,D), *i*.*e*. the expression of canonical SREBP-regulated genes (*e*.*g. HMGCR, SQLE, LDLR)* and the level of their encoded proteins remained low. A potential explanation for this discrepancy is that S1P is adequately processed, albeit to a moderately lesser degree as we observe, but fails to translocate from the ER to the Golgi where it is required for regulated proteolytic cleavage of SREBP. We evaluated this possibility by quantifying the distribution of S1P-mCherry between the ER and Golgi as measured by co-localization with Golgin97 (Figure 4E). In the absence of SPRING we found only a minimal decrease in Golgi residency of S1P (Figure 4F). Taken together, our results demonstrate that absence of SPRING has a moderate lowering effect on proteolytic maturation and cellular localization of S1P. This may contribute to abrogated SREBP-mediated signaling in cells lacking SPRING, though our results suggest that this does not represent the primary lesion.

**Figure 4.**
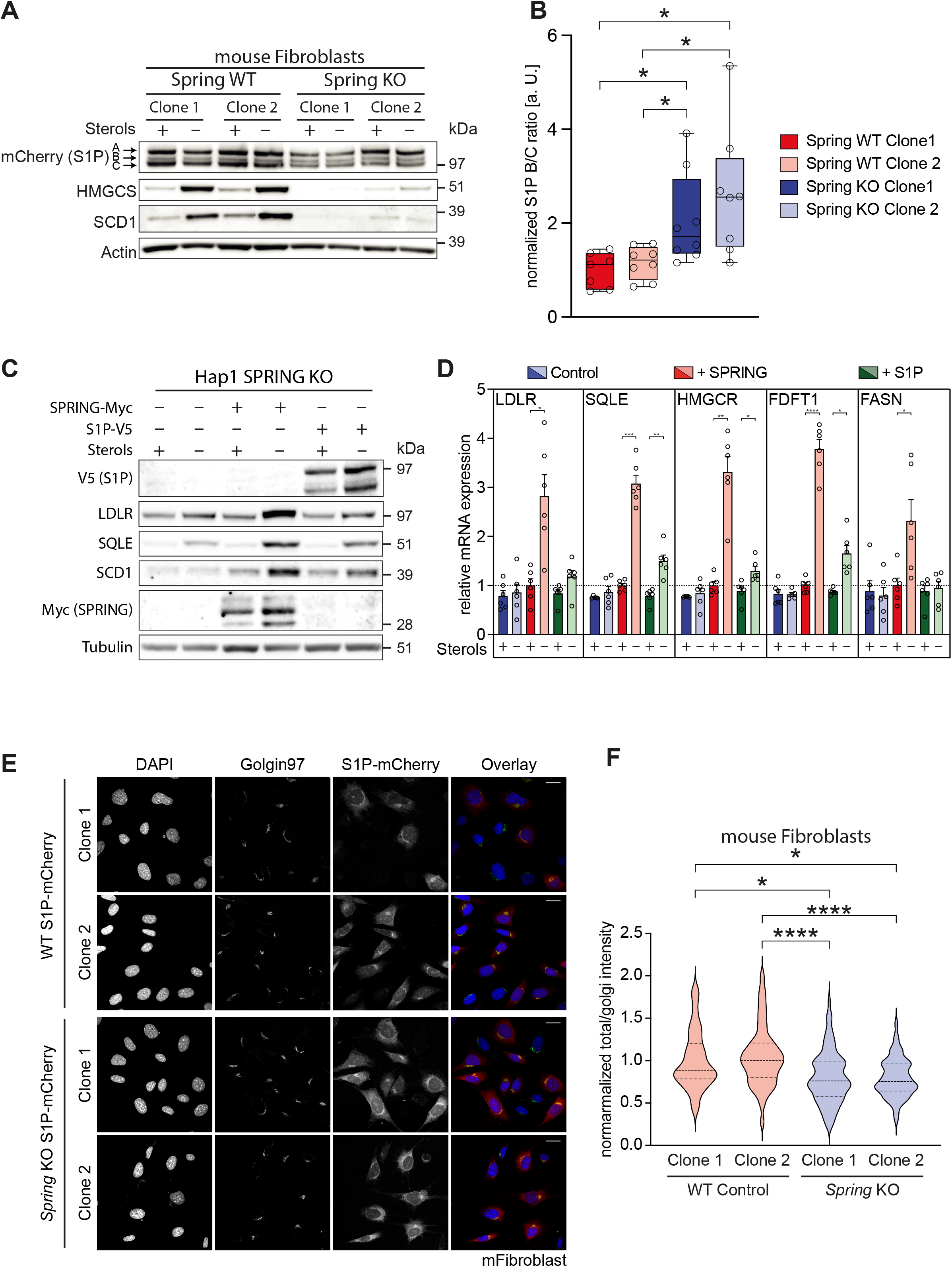
Over-expressed S1P shows a mild alteration in proteolytic processing in cells lacking SPRING and does not rescue SREBP signaling. (***A***) Control or *Spring*^*KO*^ immortalized mouse fibroblasts that stably express S1P-mCherry were grown in complete (+ sterols) or in sterol-depletion (-sterols) culture media for 16 hrs. Total cell lysates from two independent clones per genotype were immunoblotted as indicated and **(*B***) the intensity of the S1P_B_ and S1P_C_ bands was quantified. The ratio of the intensities of the S1P_B_/S1P_C_ bands is shown (N=4-5) and normalized to that of wildtype clone 1. (***C***,***D***) Hap1 SPRING^KO^ cells stably expressing the indicated constructs were grown as in (***A***), and (***C***) total cell lysates were immunoblotted as indicated, and (***D***) expression of the indicated genes was determined by qPCR (N=3). Bars and errors represent mean ± SEM. (***E***) Representative confocal images of control or *Spring*^*KO*^ immortalized mouse fibroblasts that stably express S1P-mCherry. Cells were stained with Golgin97 (Golgi) and counterstained with DAPI (nuclei). Images show individual channels and overlay. Bar 20µm (***F***) Co-localization of the mCherry (S1P) and Golgin signals was quantified in 35-110 cells from 9-12 independent images and values. Violin plots of the co-localization ratio in the different cells is shown normalized to clone 1. Median and quartiles are indicated with dashed and dotted lines, respectively. All immunoblots are representative of at least 3 independent experiments and bars and errors represent mean ± SD; * *p* < 0.05, **** *p* < 0.0001.

## Discussion

SPRING is a recently identified regulator of SREBP signaling (26-28). We have previously put forward the idea that SPRING contributes to the retrograde transport of SCAP, a necessary step for maintaining intact SREBP-dependent signaling (26). Additionally, we and others have suggested involvement of SPRING in S1P-dependent proteolytic cleavage of its targets, amongst them SREBPs, which was the focus of the current study (26,27,30). As such, the most important finding of our study is the existence of a functional interaction between S1P and SPRING that results in their mutual proteolytic processing. We further show that while S1P maturation is mildly affected when SPRING is absent, this likely does not fully explain the severe impact that the loss of SPRING has on SREBP-dependent signaling.

Several lines of evidence support a role for SPRING in S1P processing and activation. The two proteins interact, are spatially co-localized to the Golgi, and as we demonstrate here also undergo bi-directional intermolecular proteolysis which can be inhibited pharmacologically or by introducing S1P cleavage-disrupting mutations. Functionally, SPRING is needed for maximal proteolytic activation of SREBPs, ATF6, CREB3L3, and GNTAB that are all native S1P substrates (11,15). Our current study supports findings by Xiao *et al*. who also demonstrated that SPRING enhances the processing of S1P from its precursor form to the fully active C\C’ form (30). Importantly, our experiments do not distinguish whether SPRING facilitates the S1P_A->C_ or S1P_B->C_ processing. The cleavage of the B\B’ site of S1P has been proposed to occur in the ER in *cis*, while that of the C\C’ site occurs in the cis/medial Golgi. We have previously shown that targeting SPRING to the ER is unable to rescue SREBP signaling in SPRING-devoid cells (26). Hence, we reason that it is more likely that SPRING contributes to the final step of S1P proteolytic activation in the Golgi.

Intriguingly, beyond limiting proteolytic maturation of S1P, absence of SPRING also moderately attenuates its Golgi localization. The underlying reason for this remains unclear. A potential scenario explaining this may be that SPRING acts as a scaffold for assembling B/B’ cleaved S1P molecules together, and that this licenses anterograde transport of the SPRING-partially processed S1P complex to the Golgi, where low pH and higher calcium concentrations would then favor the auto-catalytic processing of S1P in *cis* for its maximal activation. Coupling this process to the S1P-mediated cleavage of SPRING may serve as a quality control step to ensure proper spatio-temporal regulation of maximal S1P activation. The proposed scaffolding model for SPRING potentially also provides an explanation as to why cleavage-resistant SPRING mutants are also able to restore SREBP-signaling in SPRING^KO^ cells; these mutants retain their ability to interact with S1P, traffic to the Golgi, and support *cis* cleavage of S1P in a manner similar to that observed when introducing cleaved SPRING construct. However, one noteworthy limitation of our studies is their reliance on (stable) over-expression of SPRING cleavage mutants. This may override stoichiometric requirements and obscure more subtle effects. Studying cells genetically engineered to endogenously produce only mutant SPRING may allow critical testing of this possibility in future studies.

We previously reported that SPRING is required for maintaining SCAP levels and localization in cells (26). The current study and that of Xiao *et al* support a role for SPRING also in S1P proteolytic maturation (30). These two processes are not interdependent, as pharmacological or genetic inhibition of SREBP-cleavage by S1P prevents retrograde transport of SCAP and promotes SCAP degradation (29). Notably, forced expression of S1P in SPRING^KO^ cells allows maturation of S1P to its C-form, albeit with reduced efficiency when compared to control cells. Nevertheless, despite attaining the fully mature S1P form, this does not restore SREBP signaling in cells lacking SPRING, in contrast to over-expression of SCAP (26). At the very least this suggests that the lesion in SREBP signaling in the absence of SPRING likely cannot be solely attributed to its role in S1P maturation. Moreover, since our studies indicate that cleavage of SPRING is not a prerequisite for SREBP signaling, the physiological significance of SPRING cleavage by S1P remains unclear. However, we note that unbiased proteomics studies of fractionated human plasma have identified multiple peptides originating solely from the cleaved SPRING fragment (*i*.*e*. post S1P cleavage site) (35-37). This supports the idea that post cleavage SPRING can be secreted into the circulation. While this may represent leakage through the secretory pathway, it is intriguing to speculate that secreted SPRING may have a role in coordinating systemic metabolic signaling.

Our understanding of the role SPRING has in lipid metabolism is largely limited to cell models. Global ablation of SPRING in mice results in embryonic lethality (26), like that seen in S1P and SCAP knockout models (38,39). The development of conditional allele models for SPRING will allow testing the tissue-specific physiological roles of SPRING and to define their overlap with that of S1P and SCAP. In summary, our study further reveals a complex role for SPRING in proteolytic maturation of S1P and in governing the SREBP pathway in cells, further highlighting the need to interrogate the role of SPRING and/or its S1P generated fragments in systemic lipid metabolism.

## Supporting information

Supp Table 1 (reagents)

Supp Table 2 (plasmids)

Supp Table 3 (antibodies)

Supp Table 4 (primers)

## Abbreviations

SREBPs: Sterol-regulatory element binding proteins
*FASN*: Fatty acid synthase
ACC: Acetyl-CoA carboxylase
HMGCR: 3-Hydroxy-3-methylglutaryl-coenzyme A reductase
SQLE: Squalene epoxidase
LDLR: Low-density lipoprotein receptor
ATF6: activating transcription factor 6
CREB: cyclic AMP-responsive element binding proteins
sSPRING: secreted SPRING
S1P: Site 1 Protease
Co-IP: Co-Immunoprecipitation

## Data availability

All data are contained within the manuscript.

## Supporting information

This article contains supporting information

## Author contributions

SH, AL, JMET, NGS and NZ concepted and designed research. SH, JMET, MV, LFZ, RO, and KN performed experiments. Data analysis and figure preparation was done by SH. Drafting, editing and revising of the manuscript was done by SH, JMET, AL and NZ.

## Acknowledgements

We thank members of the Zelcer lab and Irith Koster for their critical comments and suggestions on this study.

## Supplementary figure legends

**Supplementary Figure 1. Expression of S1P is unchanged in the absence of SPRING**. Expression of S1P was determined by qPCR in control Hap1 and Hap1-SPRING^KO^ cells (N=2). Bars and errors represent mean ± SEM

**Supplementary Figure 2. Tunicamycin-induced ATF6 cleavage is impaired in cells lacking SPRING**. Wildtype and *Spring* KO immortalized mouse fibroblasts were transfected with the indicated ATF6 or control expression constructs. Cells were treated with or without 2µg/mL tunicamycin for 6 hrs. Where indicated 25µM MG132 was added during this period to prevent the rapid proteasomal degradation of nATF6. Total cell lysates were immunoblotted as indicated. Immunoblot is representative of 3 independent experiments. pATF6, precursor ATF6; nATF6, nuclear ATF6. 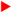 glycosylated ATF6, 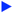 non-glycosylated ATF6 bands.

**Supplementary Figure 3. SPRING is secreted into the culture media after proteolytic cleavage by S1P**. HeLa cells were transfected with the indicated S1P and SPRING expression constructs. Total cell lysates and culture medium were collected 48 hours later and immunoblotted as indicated. Immunoblot is representative of at least 3 independent experiments.

**Supplementary Figure 4. SPRING cleavage and secretion is inhibited by introducing mutations in the S1P recognition site**. HEK293T cells were transfected with the indicated S1P and SPRING expression constructs. Total cell lysates and culture medium were immunoblotted as indicated. Immunoblot is representative of at least 3 independent experiments. uncl. SPRING; SPRING_(R43E,R45, L47)_

**Supplementary Figure 5. SPRING**_**1-44**_ **does not support SREBP signaling in Hap1-SPRING**^**KO**^ **cells**. Hap1-SPRING^KO^ cells that stably express wildtype SPRING, SPRING_1-44_, or control were cultured as described in Figure 3A-D and expression of the indicated SREBP-regulated genes was determined by qPCR. Bars and errors represent mean ± SEM; ** *p* < 0.01, *** *p* < 0.001.

**Supplementary Figure 6. Detected bands are specific for S1P-mCherry expressing cells**. Total cell lysates from control or S1P-mCherry expressing immortalized mouse fibroblasts were immunoblotted as indicated. *Note:* no signal is detected in control cells. Immunoblots are representative of at least 3 independent experiments

