## Supplemental figure 1-6 for "Intermolecular proteolytic processing of SPRING and Site-1-protease regulate SREBP signaling"

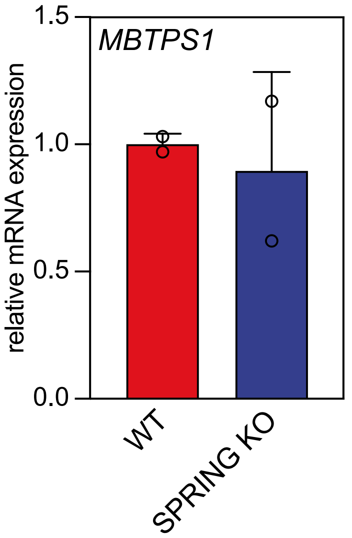

### Supplemental figure 2

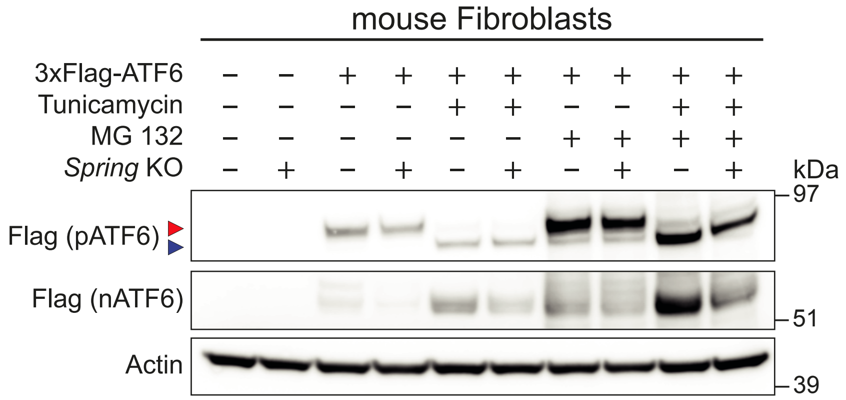

### Supplemental figure 3

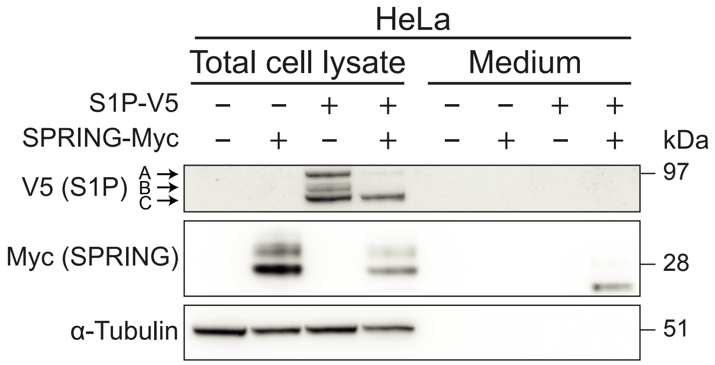

### Supplemental figure 4

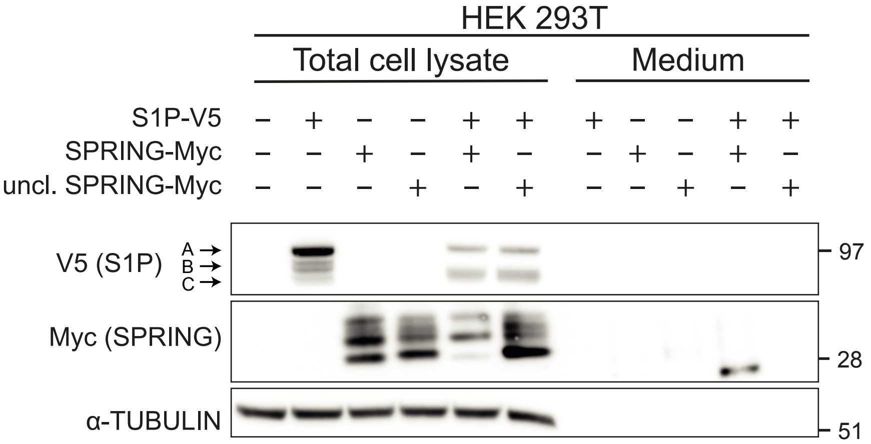

### Supplemental figure 5

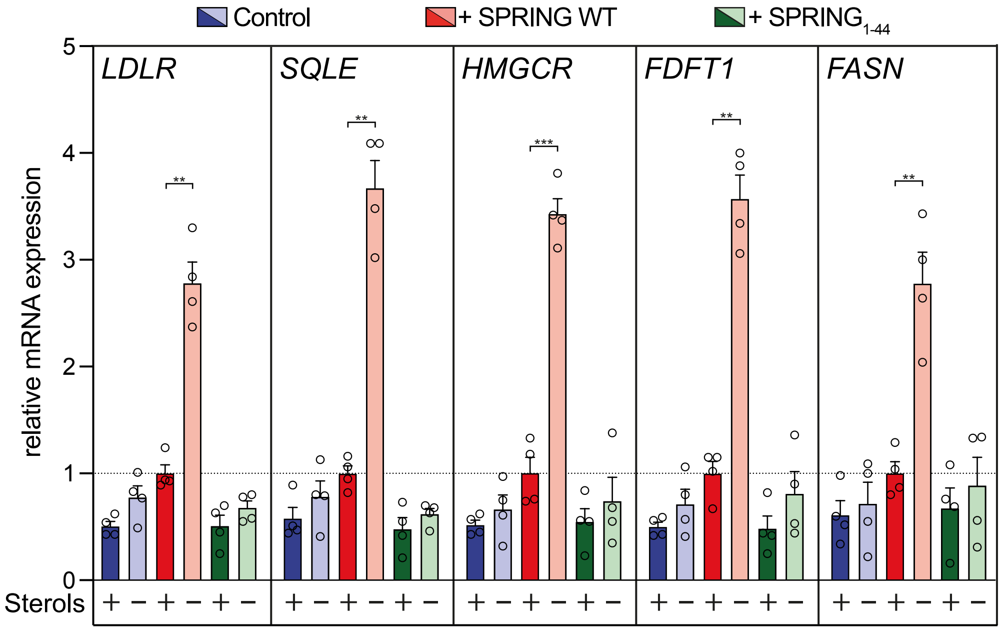

### Supplemental figure 6

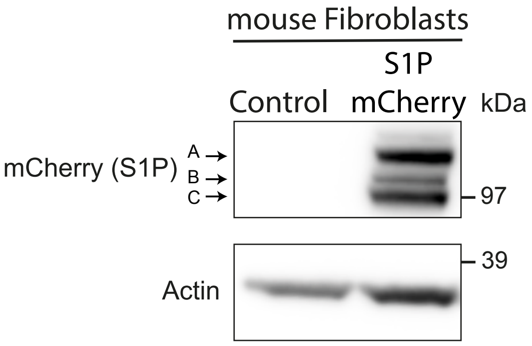
