## Supplementary material for "Intermolecular proteolytic processing of SPRING and Site-1-protease regulate SREBP signaling": Supp Table 1 (reagents)

***Supplemental table 1.*** Reagents and chemicals used in the study

| Chemicals | Source | Identifier |
| --- | --- | --- |
| Tunicamycin | Sigma | #T7765 |
| Penicillin-Streptomycin | Invitrogen | 15140-122 |
| DMEM | Invitrogen | #31966 |
| IMDM | Invitrogen | #21056-023 |
| RPMI 1640 | Invitrogen | 21875-034 |
| LPDS | Homemade | - |
| Mevalonate | Sigma | #M4667 |
| Simvastatin | Calbiochem | #567021 |
| JetPrime | Polyplus | #114-07 |
| 4% Paraformaldehyde | Sigma | F8775 |
| TripleE | Invitrogen | #12604039 |
| BP clonase | Invitrogen | #11789-020 |
| LR clonase | Invitrogen | #11791-020 |
| Poly-L-lysine | Sigma | #P9155 |
| Triton X-100 | Sigma | #T8787 |
| BSA | Sigma | #10735086001 |
| RIPA buffer | Boston Biochem | #BP-115 |
| NP-40 buffer | Boston Biochem | #BP-119 |
| Protease inhibitors | Roche | #P8340 |
| SensiFAST SYBR | Bioline | #BIO-98020 |
| MTT (3-(4,5-Dimethylthiazol-2-yl)-2,5-Diphenyltetrazolium Bromide) | Invitrogen | M6494 |
| PF 429242 dihydrochloride | bio-techne | 3354/10 |
| Mowiol | Sigma | 81381 |
| 4',6-diamidino-2-phenylindole (DAPI ) | Thermo Fisher | D21490 |
| TriReagent | Sigma | T9424 |
| Dimethyl Sulfoxide | Sigma | D8418-100ml |
| ß-Mercaptoethanol | Sigma | M3148 |
| PBS | Fresenius Gabi | M090001/02NL |
| Fibronectin | Sigma Aldrich | F0895-1MG |
