## Supplementary material for "Intermolecular proteolytic processing of SPRING and Site-1-protease regulate SREBP signaling": Supp Table 2 (plasmids)

| Construct | Source | Identifier |
| --- | --- | --- |
| pDONR221 | Thermo fisher | #12536017 |
| pLenti6.3-DEST | Thermo fisher | #V53306 |
| pLenti CMV/TO Puro Dest (1-670) | Addgene | #17293 |
| pIRES S1P-V5 EGFP | Dr. Nabil Seidah |  |
| pcDNA6.2-C-mCherry-DEST | kind gift from Dr. Paul Beare |  |
| pIRES-S1P-BTMD-EGFP | Dr. Nabil Seidah |  |
| p3xFLAG CMV 7.1 ATF6ɑ | Dr. Nabil Seidah |  |
| pmCherry-C1 | Clontech | PT3975-5 |
| pMD2.G | Addgene | #12259 |
| pMDLG/pRRE | Addgene | #12251 |
| pRSV-Rev | Addgene | #12253 |

***Supplemental table 2.*** Expression constructs and plasmids used in the study
