## Supplementary material for "Intermolecular proteolytic processing of SPRING and Site-1-protease regulate SREBP signaling": Supp Table 3 (antibodies)

***Supplemental table 3.*** Antibodies used in the study

| Antibody target | Source | Identifier | Dilution | Used for |
| --- | --- | --- | --- | --- |
| LDLR | Biovision | #3839 | 1:1000 | WB |
| SQLE | Proteintech | #12544-1-AP | 1:1000 | WB |
| ß-actin | EDM Millipore | #MAB1501 | 1:2500 | WB |
| ß-actin | Cell Signaling | 49675 | 1:2500 | WB |
| Myc-tag | Cell Signaling | #2276 | 1:1000 | WB/IP |
| V5-tag | Invitrogen | 64-0705 | 1:1000 | WB/IP |
| V5-tag | Sigma | V8137 | 1:1000 | IP |
| Flag-tag | Merck | F1804-50UG | 1:1000 | WB |
| Golgin97 | Cell Signaling | 131925 | 1:200 | IF |
| mCherry | Invitrogen | 46-0705 | 1:1000 | WB |
| Chicken anti-Rabbit IgG Alexa Fluor 488 | Invitrogen | A21441 | 1:250 | IF |
| Goat anti-Mouse IgG | Invitrogen | A28177 | 1:2500 | WB |
| Goat anti-Rabbit IgG | Invitrogen | A27036 | 1:2500 | WB |
| S1P | abcam | ab140592 | 1:500 | WB |
| alpha-Tubulin | Sigma | T9026 | 1:2000 | WB |
| HMGCS | Cell Signaling | 422015 | 1:1000 | WB |
| SCD1 | Cell Signaling | 2438S | 1:1000 | WB |
