## Supplementary material for "Intermolecular proteolytic processing of SPRING and Site-1-protease regulate SREBP signaling": Supp Table 4 (primers)

***Supplemental table 4.*** qPCR primers used in the study

| Gene | Species | Primer sequence |
| --- | --- | --- |
| LDLR fw | homo sapiens | AGGACGGCTACAGCTACCC |
| LDLR rv | homo sapiens | CTCCAGGCAGATGTTCACG |
| SQLE fw | homo sapiens | catgagtctccggaaagcag |
| SQLE rv | homo sapiens | acaacaccttcaataaactttgcat |
| HMGCR fw | homo sapiens | gacgcaacctttatatccgttt |
| HMGCR rv | homo sapiens | ttgaaagtgctttctctgtaccc |
| FDFT1 fw | homo sapiens | atgcccaagatggaccag |
| FDFT1 rv | homo sapiens | actgcgactggtctgattga |
| FASN fw | homo sapiens | caggcacacacgatggac |
| FASN rv | homo sapiens | cggagtgaatctgggttgat |
| 36B4 fw | homo sapiens | CCACGCTGCTGAACATGCT |
| 36B4 rv | homo sapiens | TCGAACACCTGCTGGATGAC |
| GAPDH fw | homo sapiens | TCTCCTCTGACTTCAACAGCGAC |
| GAPDH rv | homo sapiens | CCCTGTTGCTGTAGCCAAATTC |
